# Comparison of immunogenicity and protection efficacy of self-amplifying and circular mRNA vaccines against SARS-CoV-2

**DOI:** 10.1101/2024.08.23.609366

**Authors:** Oinam Ningthemmani Singh, Umang Berry, Garima Joshi, Tejeswara Rao Asuru, Kannan Chandrasekar, Sriram Narayanan, Puneet Srivastava, Mahima Tiwari, Souvick Chattopadhyay, Farha Mehdi, Bhisma Narayan Panda, Debasis Nayak, Shailendra Mani, Tripti Shrivastava, Gaurav Batra, C.T. Ranjith-Kumar, Prasenjit Guchhait, Milan Surjit

## Abstract

Recent advances in vaccine technology have positioned messenger RNA (mRNA) vaccines as safe and reliable options for human use. Conventionally, mRNA vaccines were designed using linear or self-amplifying mRNA (SAM), the latter considered to be superior. However, limited success was achieved with SAM vaccines during the COVID-19 pandemic. Further, studies on Circular mRNA (Circ-RNA) vaccines against the SARS-CoV-2, Ebola and monkey pox proved their efficacy. Circ-RNAs are highly stable, neither they induce inflammatory response nor require any extracellular protein for their function. Here, we compared the efficacy of SAM- and Circ-RNA vaccines using the SARS-CoV-2-RBD (receptor binding domain) as the antigen. Both SAM-RBD and Circ-RBD induced a comparable anti-RBD IgG titer and virus-neutralizing antibody titer. However, the latter induced a significantly higher memory T-cell response. Immunization with SAM- and Circ-RBD showed no mortality and improved lung pathophysiology against acute SARS-CoV-2 infection in mice. The Circ-RBD vaccine is stable for 4 weeks at 4^0^C. A bivalent vaccine containing Circ-RBD of both delta and omicron SARS-CoV-2 variants potently neutralized these viruses. These findings demonstrate Circ-RNA-RBD as an excellent vaccine candidate against COVID-19 and also provide a platform for developing bivalent Circ-RNA vaccine candidates against SARS-CoV-2 or other viruses with rapidly emerging variants.

## Introduction

Although messenger RNA (mRNA) vaccines have been researched for more than two decades, its proof of success in real-life was witnessed for the first time during the COVID-19 pandemic. Comirnaty, the first FDA (United States food and drug administration)-approved COVID-19 vaccine (developed by Pfizer-BioNTech) and Spikevax (COVID-19 vaccine developed by Moderna), both mRNA vaccines, were pivotal in controlling the COVID-19. Since then, there is considerable momentum to develop mRNA vaccines against multiple pathogens, allergic diseases and cancer (Thomas et al., 2021; Baden et al., 2021; Arevalo et al., 2022; Krienke et al., 2021; Heine et al., 2021; Parhiz et al., 2024). Advances in mRNA delivery technology, *in vitro* transcription technology and reduction in the cost of production have improved the overall yield, stability, efficacy and economic viability of the mRNA vaccine, thereby positioning it as the preferred option for vaccine development (Chaudhary et al., 2021). Both linear mRNA (such as Comirnaty and Spikevax) and self-amplifying mRNA (SAM) (such as Gemcovac)-based vaccines have been approved for use in human (Thomas et al., 2021; Baden et al., 2021; Saraf et al., 2024). More recently, several independent studies have demonstrated the efficacy of circular mRNA (Circ-RNA) vaccines (Qu et al., 2022; Amaya et al., 2023; Seephetdee et al., 2022; Zhou et al., 2024).

Linear mRNA-encoding the vaccine antigen is generally small in size (compared to the SAM), contain a methyl-guanosine cap analog at the 5’-end, poly A-tailed at the 3’-end and the antigen-encoding sequence is flanked by untranslated regions (UTR) at both sides. A major challenge of linear mRNA-based vaccines is low stability of the input mRNA. Stability of the linear mRNAs are increased by incorporating pseudo-uridine or N6-Methyladenosine (m6A) into the sequence (Karikó et al., 2008; Wang et al. 2014). SAM vaccines are generally based on engineered genome of alphaviruses such as the Venezuela Equine Encephalitis virus (VEEV), Semliki Forest Virus (SFV) or Sindbis virus (SINV) (Bloom et al., 2020). Here, the viral structural protein-encoding sequence is replaced with the sequence of the vaccine antigen so that non-structural proteins of the virus enables replication the viral genome without production of infectious progeny virions. Thus, SAM vaccines should produce the required quantity of antigen for a prolonged period with less amount of the input RNA (compared to the linear mRNA), leading to a stronger and more durable immune response (Schmidt and Schnierle, 2023). However, use of viral replication machinery for amplifying the input RNA has certain disadvantages such as: size of the SAM RNA is considerably longer as it contains the sequence to express viral non-structural proteins; viral RNA-dependent RNA polymerase, which is required for replication of the SAM RNA has low fidelity, increasing the possibility of introducing mutation(s) in the antigen-encoding sequence; pre-existing anti-vector immunity remains a concern; generation of dsRNA during the replication of SAM might activate host defence machinery leading to degradation of SAM; possibility of uncontrolled and excessive replication of SAM due to mutations in the non-structural proteins cannot be ruled out. Notably, one of the SAM vaccines (COVAC1) failed to induce 100% seroconversion while another SAM vaccine (ARCT154) showed a very good response in phase I clinical trials (Pollock et al., 2022; Oda et al., 2024). Gemcovac (Gennova), a commercially available SAM vaccine, also shows very good protection efficacy in clinical trials (Saraf et al., 2024).

Circular RNAs (Circ-RNA) are covalently closed single-stranded RNA without free 5’- and 3’- ends. It is produced naturally through back-splicing, which involves the connection of the 3’- end of an exon to the 5’- end of another exon, creating a close loop RNA. The circular structure provides resistance to most of the RNases except RNase A, RNase T1 and RNase T2, thus significantly increasing their half-life compared to linear mRNAs. Endogenous Circ-RNAs act as miRNA sponges to regulate various cellular processes by binding to nucleic acids and proteins, thereby modulating transcription and cellular signalling pathways (Liu and Chen, 2022). Circ-RNAs do not induce the cellular innate immune pathway RNA sensors (Wesselhoeft et al., 2019; Breuer et al., 2022). Recent reports have demonstrated the ability of engineered Circ-RNAs to express the vaccine antigens using internal ribosomal entry site (IRES)-mediated translation, thereby eliminating the requirement of capping of the 5’-end of the mRNA (Wesselhoeft et al., 2018a; Qu et al., 2022). Circ-RNA-based vaccines against the Severe acute respiratory syndrome-coronavirus-2 (SARS-CoV-2) and Monkey pox has been shown to induce potent immune response and protect the animals from challenge with the respective viruses (Qu et al., 2022; Zhou et al., 2024). Notably, in the case of SARS-CoV-2, Circ-RNA immunization produced a higher T_H_1 biased humoral immune response and a higher neutralizing to binding antibody ratio than its mRNA counterpart, supporting its protection efficacy and superior potential for evading the antibody-dependent enhancement (ADE) (Qu et al., 2022).

SARS-CoV-2, the causative agent of the coronavirus-induced disease-19 (COVID-19), is responsible for the worst global pandemic of the 21^st^ century. So far approximately 776 million COVID-19 cases and 7 million deaths have been reported globally (https://data.who.int/dashboards/covid19). The viral genome shows high mutation rate, leading to emergence of multiple variants, some of which, such as the B.1.617.2 (Delta) and the Omicron B.1.1.529 (BA.1) variants pose serious health issues (Tao et al., 2021; Carabelli et al., 2023). Omicron B.1.1.529 (BA.1) became the most dominant circulating strain with 22 subvariants accounting for 87% of infections. Omicron strain dominated the previous variant of concern (VOC) in 2022 and carries the most significant number of mutations than the previous VOC, with 37 mutations within its spike protein. 15 mutations at the RBD region covering almost all the critical mutation sites, such as K417N, S477N, T478K, E484A and N501Y make more transmissible and immune evasion to pre-existing immunity. Previous non-Omicron infection or vaccination has little protection against the Omicron variant (Cameroni et al., 2021; Liu et al., 2022; Planas et al., 2022). The Omicron sub-variants, including the BA.4/5 and XBB, continue to emerge, with total immune escaping potential to the previous vaccination. Few studies have shown the broader protective efficacy of bivalent mRNA and nanoparticle vaccines composed of a mixture of antigens from the SARS-CoV-2-Beta and Delta or Delta and Omicron variants (Yuan et al., 2022; Corleis et al., 2023; Schmidt and Schnierle, 2023). Despite development of several vaccines against the SARS-CoV-2, there is limited access to vaccines globally and there is scope for improving the efficacy and duration of protection offered by the vaccines.

Being the viral outer surface protein, spike protein of the SARS-CoV-2 is the main target for vaccine development. The receptor binding domain (RBD) of the spike protein, which accounts for one-fifth of the total protein, produced over 90% of the neutralizing antibody in infected patients (Piccoli et al., 2020). There are no reports of antibody dependent enhancement (ADE) by vaccines based on the full length spike protein of the SARS-CoV-2. However, possibility of ADE induction by the spike protein of emerging SARS-CoV-2 variants cannot be ruled out. Therefore, RBD would be the best target for developing SARS-CoV-2 vaccines.

In the present study, we compared the immunogenicity and virus neutralization potential of the Circ-RNA and SAM-RNA vaccine candidates using the SARS-CoV-2-RBD as the vaccine antigen. Although both Circ-RBD (Circ-RNA backbone) and SAM-RBD (SAM-RNA backbone) elicited comparable humoral and neutralizing antibody response, the former induced a significantly higher T_H_1-biased cellular immune response and protected mice from challenge with the SARS-CoV-2. Further, Circ-RBD vaccine was stable in refrigerator, induced a durable neutralizing antibody response and a bivalent vaccine formulation encoding the RBD of Delta (Circ-RBD^Del^) and Omicron (Circ-RBD^Omi^) variants induced high cross-neutralizing antibody response against both variants, confirming the protective efficacy of the Circ-RNA vaccines. Significance of these findings in designing better mRNA vaccines based on circular RNA technology is discussed.

## Results

### Production and characterization of the SARS-CoV-2 SAM-RBD and Circ-RBD vaccine formulations

Self-splicing property of the Anabaena intron (Group 1) was utilized to generate the Circ-RNA encoding the SARS-CoV-2-RBD (Wuhan isolate) fused to the CD5 signal sequence (SS) and the sequence encoding the trimerization motif of bacteriophage T4 fibritin (foldon) at the N- and C-terminus, respectively. The fusion protein was translated under control of a Coxsackie virus B3 (CVB3) internal ribosome entry site, as reported earlier (Wesselhoeft et al., 2018a). A bacteriophage T7 promoter sequence was placed upstream of the anabaena intron sequence at the 5’-end to enable *in-vitro* transcription-mediated synthesis of Circ-RBD. A Semliki Forest virus (SFV) replicon was engineered to produce the self-amplifying RNA encoding the SARS-CoV-2-RBD (SAM-RBD). RBD-encoding sequence fused to the CD5 signal sequence and T4 fibritin foldon sequence at its 5’- and 3’-ends was inserted into the SFV replicon. *In-vitro* transcription was carried out using the SP6 polymerase, followed by Cap1-capping of the RNA. Sized integrity of the RNAs were verified by spectrophotometry and formaldehyde-agarose gel electrophoresis. Expected sized of the RNAs were observed (Figure 1A). An aliquot of the RNAs were treated with the RNase R to check their circularization status. SAM-RBD was completely digested by RNase R treatment whereas majority of Circ-RBD was resistant, confirming its circularization (Figure 1A). Next, RNA was encapsulated into the lipid-nanoparticle (LNP) by microfluidic mixing and the diameter of the resulting RNA-LNP formulation was measured. The average diameter of the SAM-RBD-LNP and the Circ-RBD-LNP was found to be 140 and 136nm, respectively (Figure 1B). Average polydispersity index (PDI) of the SAM-RBD-LNP and the Circ-RBD-LNP was 0.06 and 0.04, respectively (Figure 1C). LNP encapsulation and stability of the SAM-RBD and the Circ-RBD RNA in the LNP complex was verified by resolving the RNA-LNP complex with or without treatment with Triton X-100 (disrupts the LNP). No RNA was visible in the SAM-RBD-LNP and the Circ-RBD-LNP complex-containing samples (Figure 1D). Treatment of the sample with Triton X-100 results in detection of the SAM-RBD and Circ-RBD (Figure 1D). No difference was observed in the quality of the SAM-RBD and Circ-RBD RNA before and after LNP encapsulation (Figure 1D). Percentage LNP encapsulation of the SAM-RBD and Circ-RBD RNA was quantified using Ribogreen. 98.3% and 95.9% of SAM-RBD and Circ-RBD RNA, respectively, was LNP-encapsulated (Figure 1E). Scanning electron microscopy analysis of the SAM-RBD-LNP and Circ-RBD-LNP complexes revealed spherical structure with average diameter of 136nm and 149nm respectively (Figure 1F). Next, SAM-RBD-LNP and Circ-RBD-LNP complexes were transfected into HEK293T cells to detect the expression of RBD. Western Blot analysis of the whole cell extract using the anti-RBD antibody confirmed expression of the RBD protein (Figure 1G, upper panel). An aliquot of the whole cell extract was probed using the anti-GAPDH antibody to check equal loading of the sample (Figure 1G, lower panel). Secretion of RBD protein into the culture medium was checked by anti-RBD western blot of the culture medium collected from the cells used in the above experiment (shown in Figure 1G). RBD was detected in the culture medium as 2 distinct bands, likely representing the glycosylated variants of the same protein (Figure 1H). Aliquots of the same sample were resolved in non-reduced state to check trimerization status of the RBD protein. As expected, anti-RBD western blot detected a band of approximately 100kDa in the non-reduced samples, indicative of RBD-trimer (Figure 1I). Possible activation of the innate immune pathways by the SAM-RBD and Circ-RBD RNAs were checked by RT-qPCR analysis of IFN-β, RIG-I, OASL, PKR and OAS1 genes. None of these genes were induced by either of the RNAs (Figure 1J). As a positive control, cells were transfected with poly I:C in parallel, which is known to induce the cellular antiviral response pathway (Figure 1J). Collectively, these results confirmed LNP encapsulation of SAM-RBD and Circ-RBD RNA and expression of RBD protein trimer by both the RNAs. Importantly, western blot analysis also demonstrated that approximately same amount of RBD was produced by both the SAM-RBD and the Circ-RBD. Therefore, our experimental condition was suitable to compare the immune response induced by the SAM-RBD and the Circ-RBD.

**Figure 1.**
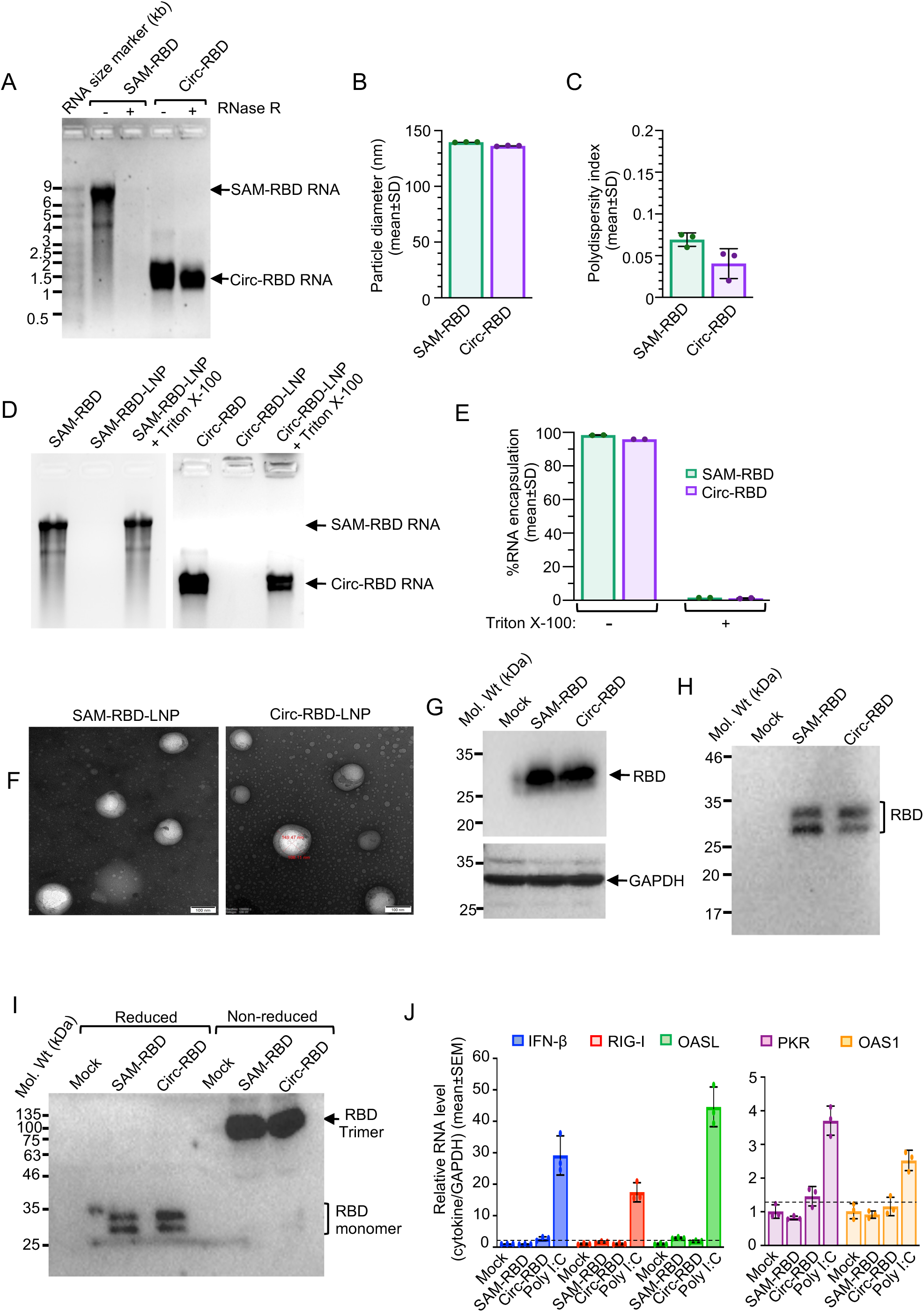
Production and characterization of the self-amplifying and circular RNA vaccine candidates. (A) Agarose gel electrophoresis analysis of *in vitro* transcribed and purified SAM-RBD and Circ-RBD RNA with (+) and without (-) treatment with RNase R. (B) Average particle size of SAM-RBD-RNA-LNP and Circ-RBD-RNA-LNP complex. (C) Average polydispersity Index of SAM-RBD-RNA-LNP and Circ-RBD-RNA-LNP complex. (D) Agarose gel electrophoresis analysis of SAM-RBD and Circ-RBD-RNA incorporation into the LNP on RNA-LNP formulation treated with Triton X-100, as indicated. (E) Average percent SAM-RBD and Circ-RBD RNA incorporation into the LNP on RNA-LNP formulation treated with Triton X-100, as indicated. (F) Scanning electron microscopy image of SAM-RBD-LNP and Circ-RBD LNP. Scale:100nm. (G) Western blot analysis of RBD protein in HEK239T cells transfected with SAM-RBD and Circ-RBD RNA (upper panel). Mock: Lysate from HEK293T cells treated with LNP. Lower panel: Anti GAPDH western blot of aliquots of cell lysate used in the upper panel. (H) Anti-RBD western blot of RBD protein in the cell culture medium of HEK239T cells mentioned in (G). (I) Anti-RBD western blot of aliquots of sample used in (H) under reducing and non-reducing conditions, as indicated. (J) RT-qPCR analysis of the indicated genes in total RNA isolated from HEK239T cells transfected with SAM-RBD or Circ-RBD-RNA, as indicated. Mock: lipofectamine 3000 transfected cells. A set of samples from poly I:C transfected cells were simultaneously processed, as positive control. Dotted line indicates the background level. Data are mean±SD of three independent experiments. *P*-values were calculated by the non-parametric student’s t-test, using the Mann-Whitney test.

### SAM-RBD and Circ-RBD RNA vaccines induced significant T_H_1-biased humoral and cellular immune response in mice

To select the injection route and the optimal RNA dose, Balb/c mice were immunized via intradermal (ID) or intramuscular (IM) route with freshly prepared vaccine formulation containing 25μg Circ-RBD RNA. Two weeks later, immunization was repeated and blood was collected after four weeks (day 28). Both ID and IM routes showed similar anti-RBD IgG titer (average values: 6×10^5^ and 1×10^6^, respectively) and virus neutralization titer (average NT_50_= 1.7×10^3^ and 1.9×10^3^, respectively) (Figures S1A and B). ID immunization route was selected for all subsequent experiments. Next, mice were injected with 5, 10 and 25μg of Circ-RBD-RNA containing vaccine formulations in a two-dose immunization schedule as described above. Measurement of anti-RBD IgG titer revealed average values to be 2.2×10^4^, 2.8×10^5^, 6×10^5^ and average virus neutralization titer (NT_50_) was found to be 0.5×10^3^, 1.5×10^3^, 0.8 ×10^3^, in 5, 10 and 20μg RNA containing formulations, respectively (Figures S1C and S1D). There was no significant difference in values obtained between all 3 doses, hence 10μg RNA containing formulation was selected for all subsequent experiments.

In order to evaluate the immune response generated by the Circ-RBD and SAM-RBD, 10μg of each RNA containing vaccine formulation was intradermally injected into Balb/c mice in two dose immunization schedules at two-week interval, followed by collection of blood and spleen samples for analysis of different immune parameters (Figure 2A). Significant level of anti-RBD IgM was detected in the sera of mice immunized with both SAM-RBD and Circ-RBD RNA (Figure 2B). Importantly, no significant difference was observed in the level of IgM between SAM-RBD and Circ-RBD immunized mice (Figure 2B). Measurement of anti-RBD IgG titer and virus neutralization titer (NT_50_) in the sera collected on day 28 from SAM-RBD and Circ-RBD immunized mice revealed the average values to be 2.8×10^5^ and 3.2×10^5^ (anti-RBD-IgG titer) and 1×10^3^ and 1.5×10^3^ (NT_50_), respectively (Figures 2C and 2D). IgG-isotyping of aliquots of the same sera showed comparable values between SAM-RBD and Circ-RBD groups (Figures 2E-2G). The ratio of anti-RBD (IgG2a+IgG2b) to IgG1 was found to be 1.9 and 1.8, respectively, indicating a T_H_1 response (Figure 2H). Next, splenocytes from these mice were cultured *in vitro* and induced with overlapping peptide pool corresponding to the RBD region, followed by selection of CD4^+^ and CD8^+^ T cells and quantification of intracellular T_H_1 (IFN-γ, TNF-α) and T_H_2 (IL-4) cytokines. Both CD4^+^ and CD8^+^ T cells of mice immunized with Circ-RBD induced significant IFN-γ and TNF-α response (Figures 2I-2L). A significant IFN-γ and TNF-α response was observed in CD4^+^ and CD8^+^ T cells, respectively, in SAM-RBD immunized mice (Figures 2I and 2L). An increased trend was observed in IFN-γ and TNF-α level in CD8^+^ and CD4^+^ T cells, respectively, in SAM-RBD immunized mice although it was not statistically significant (Figures 2J and 2K). There were no significant changes in IL-4 response in both SAM-RBD and Circ-RBD immunized mice, indicating that both vaccine formulations induced T_H_1-biased T-cell immune response (Figures 2M and 2N). Representative FACS plots are shown (Figure S2). Collectively, these results demonstrate that both SAM-RBD and Circ-RBD induced a robust and comparable humoral immune response, however, the latter induced a better T_H_1-biased cell-mediated immune response.

**Figure 2.**
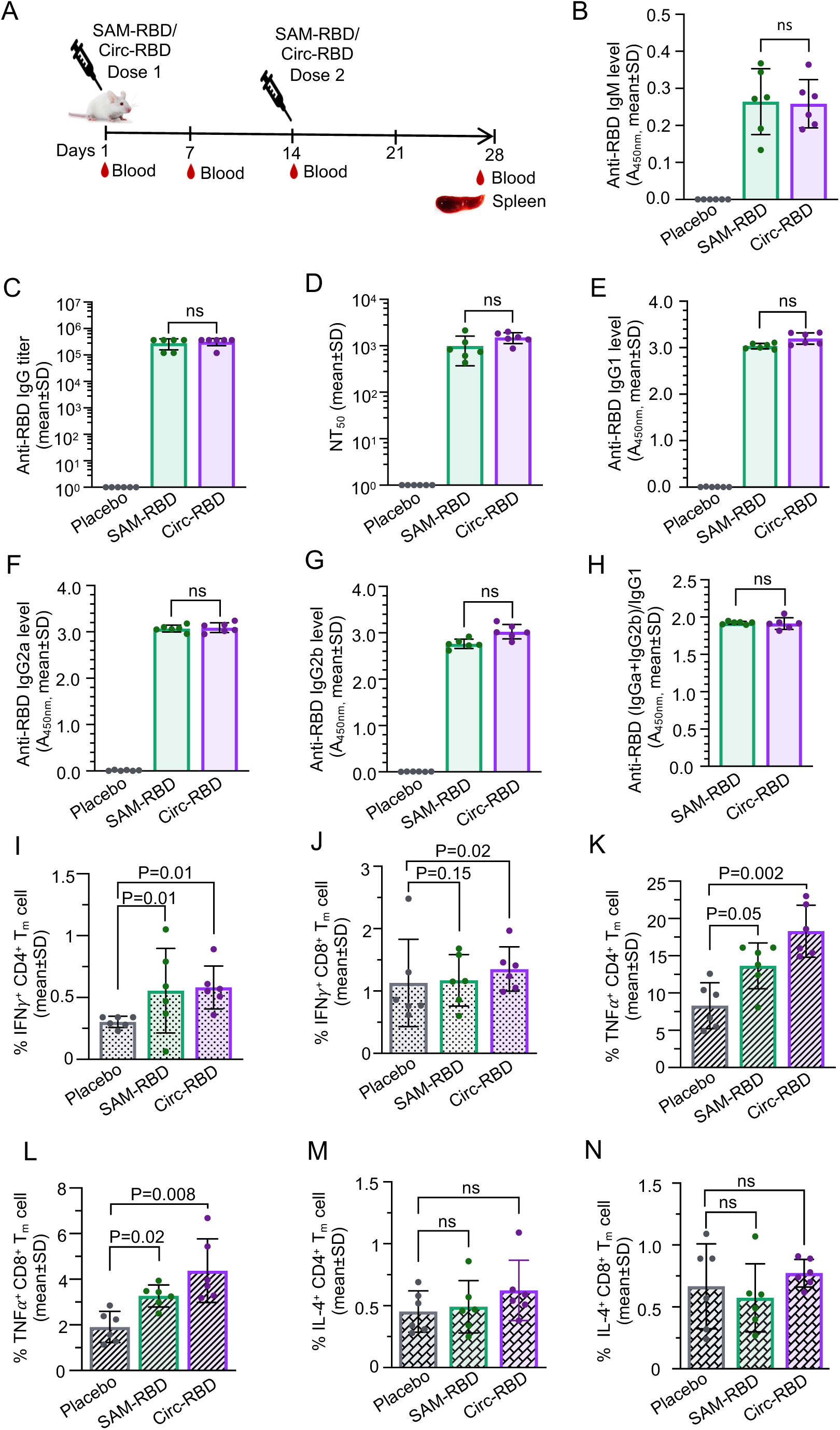
Immunogenicity assessment of SAM-RBD and Circ-RBD-RNA vaccine formulations. (A) Schematic of the immunization study. Balb/c mice were intradermally injected with 10μg SAM-RBD or Circ-RBD-RNA-LNP on day 1 and 14. Blood was collected via retro-orbital route prior to the first immunization and subsequently on day 7 and day 28. Animals were euthanized on day 28, blood and spleen were collected. (B) Anti-RBD IgM level at 1 week post-immunization in the sera (1:500 dilution) of mice injected with the placebo (PBS), SAM-RBD and Circ-RBD-LNP, evaluated by ELISA. (C) Anti-RBD IgG titre at 4 weeks post-immunization in the sera of mice described in (B), evaluated by ELISA. (D) 50% virus neutralization titre (NT_50_) at 4 weeks post-immunization in the sera of mice described in (B), evaluated by PRNT assay. (E) Anti-RBD IgG1 level (1:500 dilution) at 4 weeks post-immunization in the sera of mice described in (B), evaluated by ELISA. (F) Anti-RBD IgG2a level (1:500 dilution) at 4 weeks post-immunization in the sera of mice described in (B), evaluated by ELISA. (G) Anti-RBD IgG2b level (1:500 dilution) at 4 weeks post-immunization in the sera of mice described in (B), evaluated by ELISA. (H) Ratio of anti-RBD T_H_1 antibody (IgG2a and IgG2b) to T_H_2 (IgG1) antibody in the sera of mice injected with the SAM-RBD and Circ-RBD at 4 weeks, evaluated by ELISA. (I) Percentage of IFNγ and CD4 positive T-cells in the splenocytes, following stimulation with the SARS-CoV-2-Spike-RBD peptide pool, evaluated by FACS. (J) Percentage of IFNγ and CD8 positive T-cells in the splenocytes, following stimulation with the SARS-CoV-2-Spike-RBD peptide pool, evaluated by FACS. (K) Percentage of TNFα and CD4 positive T-cells in the splenocytes, following stimulation with the SARS-CoV-2-Spike-RBD peptide pool, evaluated by FACS. (L) Percentage of TNFα and CD8 positive T-cells in the splenocytes, following stimulation with the SARS-CoV-2-Spike-RBD peptide pool, evaluated by FACS. (M) Percentage of IL-4 and CD4 positive T-cells in the splenocytes, following stimulation with the SARS-CoV-2-Spike-RBD peptide pool, evaluated by FACS. (N) Percentage of IL-4 and CD8 positive T-cells in the splenocytes, following stimulation with the SARS-CoV-2-Spike-RBD peptide pool, evaluated by FACS. In all images, bars represent the average and dots represent the value obtained from each animal. Data are mean±SD, *p*-values were calculated by the non-parametric student’s t-test, using the Mann-Whitney test (B-H) and the non-parametric one-way ANOVA, using the Kruskal-wallis test (I-N). Values showing distinct difference from the majority in the group were not considered for calculating the *p*-value in the panels I-L.

### Immunization with the SAM-RBD and Circ-RBD RNA vaccines protect mice from the SARS-CoV-2 infection and prevents COVID-19 associated pathological changes

In order to evaluate the protection efficacy of the Circ-RBD vaccine, we established a mouse model of the SARS-CoV-2 infection that produces a severe disease phenotype. Intratracheal administration of the replication deficient adenovirus expressing the human angiotensin converting enzyme 2 (Ad5CMV-hACE2, denoted as hACE2 hereafter) has been shown to express the hACE2 in mouse lungs within 5 days of transduction (Gu et al., 2021). We hypothesized that increased hACE2 expression in the lungs will permit more pronounced SARS-CoV-2 infection in the lungs, leading to a more severe disease phenotype. To test the hypothesis, mice were transduced with the hACE2 via intratracheal administration, followed by intranasal infection with the 10^5^ PFU of SARS-CoV-2 (Wuhan isolate), as described (Hassan et al., 2020). Body weight of the animals were measured every day, blood was collected prior to infection and upto 3 weeks after infection. Survivability of the animals were monitored. Few animals were sacrificed each week and blood and lung tissue were collected for analysis. Results obtained in the pilot experiment confirmed that our animal model displayed a more severe pathogenic effect, including lethality in some animals. Protection efficacy of the SAM-RBD and Circ-RBD RNA vaccines were compared in the above model. Mice were immunized twice at 2 weeks intervals, followed by intratracheal transduction of the hACE2 (on day 28). 10^5^ PFU of the SARS-CoV-2 (Wuhan) was intranasally administered after 5 days and disease parameters were followed up for next three weeks (up to day 54). The experimental plan is summarized (Figure 3A). In the placebo group, there was a marked reduction in the body weight from the 2^nd^ day of infection up to the 7^th^ day, after which there was a gradual increase in the body weight up to the 21^st^ day (Figure 3B). In the case of SAM-RBD and Circ-RBD vaccine immunized group, weight loss was observed between the 2^nd^ to 4^th^ day post-infection after which the animals gained weight, indicating that both vaccines protected the animals from progressing to severe disease pathology (Figure 3B). Placebo group animals appeared sick, displayed hunchback posture, barely moved and importantly, three animals died during the observation period (one animal each died on day 36, 38 and 40) (Figure 3C). The remaining animals started to gain weight from day 41 onwards. Notably, SAM-RBD and Circ-RBD vaccine immunized animals were almost normal throughout the observation period.

**Figure 3.**
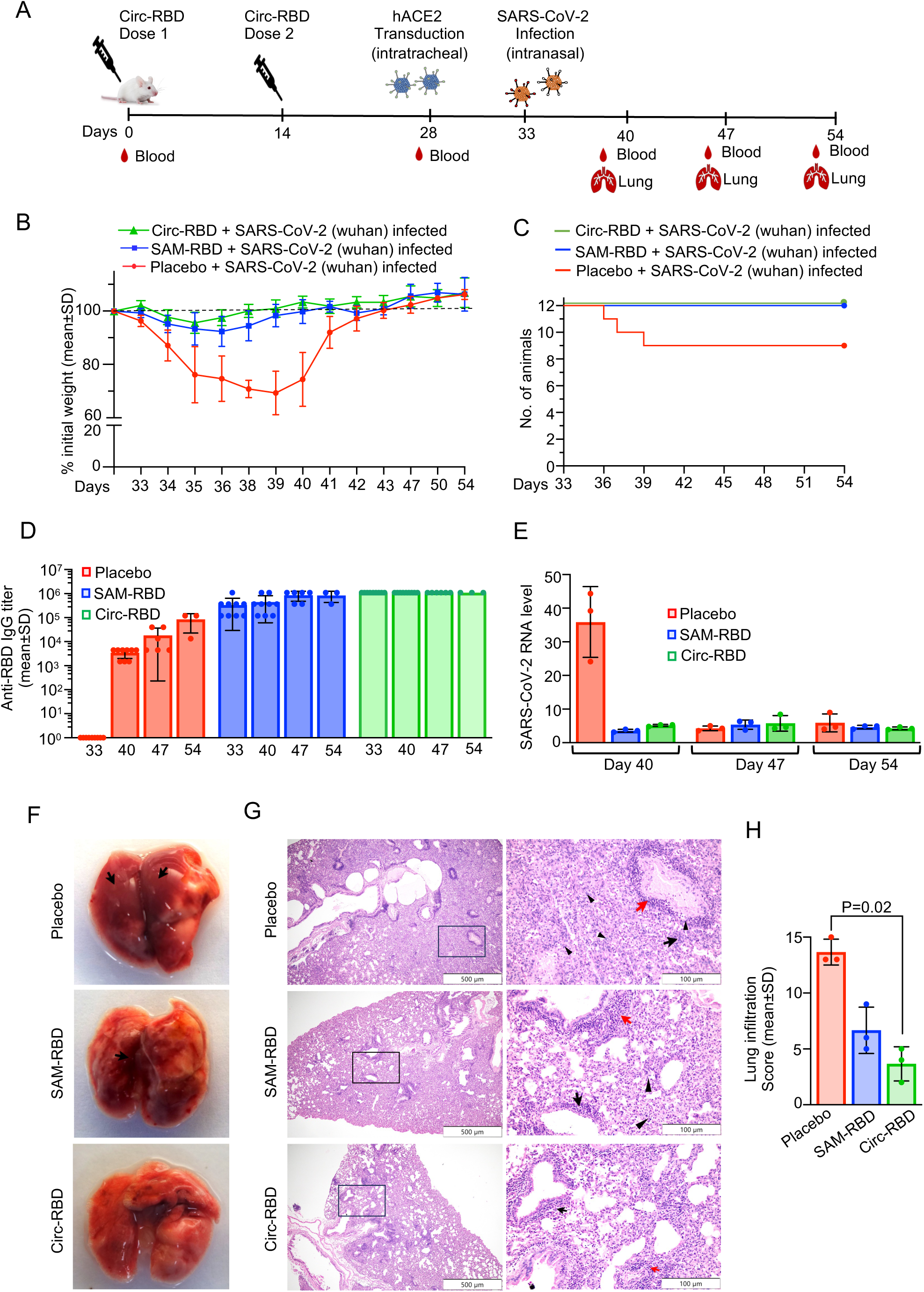
Immunization with SAM-RBD and Circ-RBD RNA vaccines protect mice from intratracheal SARS-CoV-2 challenge-induced disease symptoms. (A) Schematic of immunization and challenge study. Blood was collected before immunization on day 0 and day 14. Intratracheal transduction of hACE2-adenovirus was done on day 28, followed by intranasal infection with SAR-CoV-2 (Wuhan) on day 33. Mouse weight was measured every day from day 33-54. Mice were sacrificed (3 animals at each time point) on day 40, 47 and 54 and blood and lung tissues were collected. (B) Weight of each mouse in placebo, SAM-RBD and Circ-RBD RNA vaccine immunization group was measured every day from day 33-54 and the percent weight change with respect to the weight on day 33 (considered 100%) is plotted in the graph. (C) Survival of mice in placebo, SAM-RBD and Circ-RBD RNA vaccine immunized groups were monitored every day from day 33-54. (D) Anti-RBD IgG titer in the sera of placebo, SAM-RBD and Circ-RBD RNA vaccine immunized, SARS-CoV-2 (Wuhan) infected mice as indicated in (A). Sera collected on day 40, 47 and 54 were evaluated by ELISA. (E) Measurement of viral RNA level in aliquots of the lung tissue of placebo, SAM-RBD and Circ-RBD RNA vaccine immunized, SARS-CoV-2 (Wuhan) infected mice as indicated in (A). Viral RNA level was determined by RT-qPCR in samples collected on day 40, 47 and 54. (F) Lung morphology of the placebo, SAM-RBD and Circ-RBD RNA vaccine immunized, SARS-CoV-2 (Wuhan) infected mice, collected on day 40. (G) Representative H&E stained image of lung tissue section prepared from placebo, SAM-RBD or Circ-RBD RNA vaccine immunized, SARS-CoV-2 (Wuhan) infected mice, collected on day 40. Image were acquired at 40X (left panel) and 200X magnification (right panel). Black and red arrows indicate peribronchiolar and perialveolar infiltration, respectively. Black arrow-head indicates neutrophil infiltration. (H) Lung infiltration score of placebo, SAM-RBD and Circ-RBD RNA vaccine immunized, SARS-CoV-2 (Wuhan) infected mice, evaluated by analysis of H&E stained images of tissue sections prepared from lungs collected on day 40. In (B), (D), (E) and (H), data are shown as the mean±SD and significance was determined using unpaired student’s t-test, Mann-Whitney test.

Anti-RBD IgG titer was measured in the placebo and vaccinated groups on day 33, 40, 47 and 54. There was no anti-RBD IgG in the placebo group on day 33 and an average titer of 5*10^3^ was observed after one week of infection, which increased upto 10^5^ by day 54 (Figure 3D). SAM-RBD vaccine immunized group showed an average anti-RBD IgG titer of 10^5^-10^6^ (Figure 3D). Estimation of viral RNA level and viral load in the samples collected between 33-54 days whereas Circ-RBD vaccine immunized group showed in the lung revealed higher viral RNA level (approximately ten-fold higher) and higher viral load (10^3^ PFU/mg of lung) in the lung tissue of placebo group (samples collected on day 40) compared to the SAM-RBD and Circ-RBD vaccine immunized groups (Figures 3E, S3A). Both viral RNA level and viral load were significantly less in the placebo group samples collected on day 47 and 54, which inversely correlated with the anti-RBD IgG titer observed at the same time point (Figures 3E, S3A). Morphology and histopathology analysis of lung tissue of placebo group collected on day 40 showed multiple lesions, pneumonia and infiltration of inflammatory cells whereas SAM-RBD and Circ-RBD vaccine immunized lung samples were significantly better (Figures 3F, 3G). A similar profile was observed in the lung samples collected on day 47 and 54 (Figures S3B, S3C). Quantification of lung infiltration score in the slides further confirmed the above observation (Figure 3H). Collectively, these finding confirmed the functional ability of the SAM-RBD and the Circ-RBD vaccines in protecting the mice against SARS-CoV-2 infection.

### Circ-RBD vaccine formulation is stable in refrigerator (4^0^C) and induces a durable immune response in both female and male mice

Since Circ-RBD vaccine induced a better overall immune response than the SAM-RBD, we evaluated the stability and durability of the immune response induced by it. To evaluate the effect of prolonged storage (up to 4 weeks) of the Circ-RBD vaccine in the refrigerator (4^0^C), it was stored in aliquots in the refrigerator for 2 or 4 weeks, followed by evaluation of its physical and functional characteristics. No change in particle diameter, polydispersity index and percent RNA encapsulation was observed between the samples stored for 2 and 4 weeks, compared to the freshly prepared sample (0 week) (Figures 4A-4C). Mice were injected with aliquots of the same samples in a two-dose immunization schedule (as described above), followed by collection of blood on the 28^th^ day. There was no difference in the anti-RBD IgG titer and the virus neutralization titer (NT_50_) between freshly prepared and two weeks-stored samples (IgG titer=2.8×10^5^ and 4×10^5^, NT_50_=1.7×10^3^ and 1.3×10^3^ respectively) (Figures 4D and 4E). There was approximately one log drop in the anti-RBD IgG titer and a 1.5 log drop in the virus neutralization titer in the samples stored for 4 weeks at 4^0^C (IgG titer=0.6×10^5^, NT_50_=0.04×10^3^) (Figures 4D and 4E).

**Figure 4.**
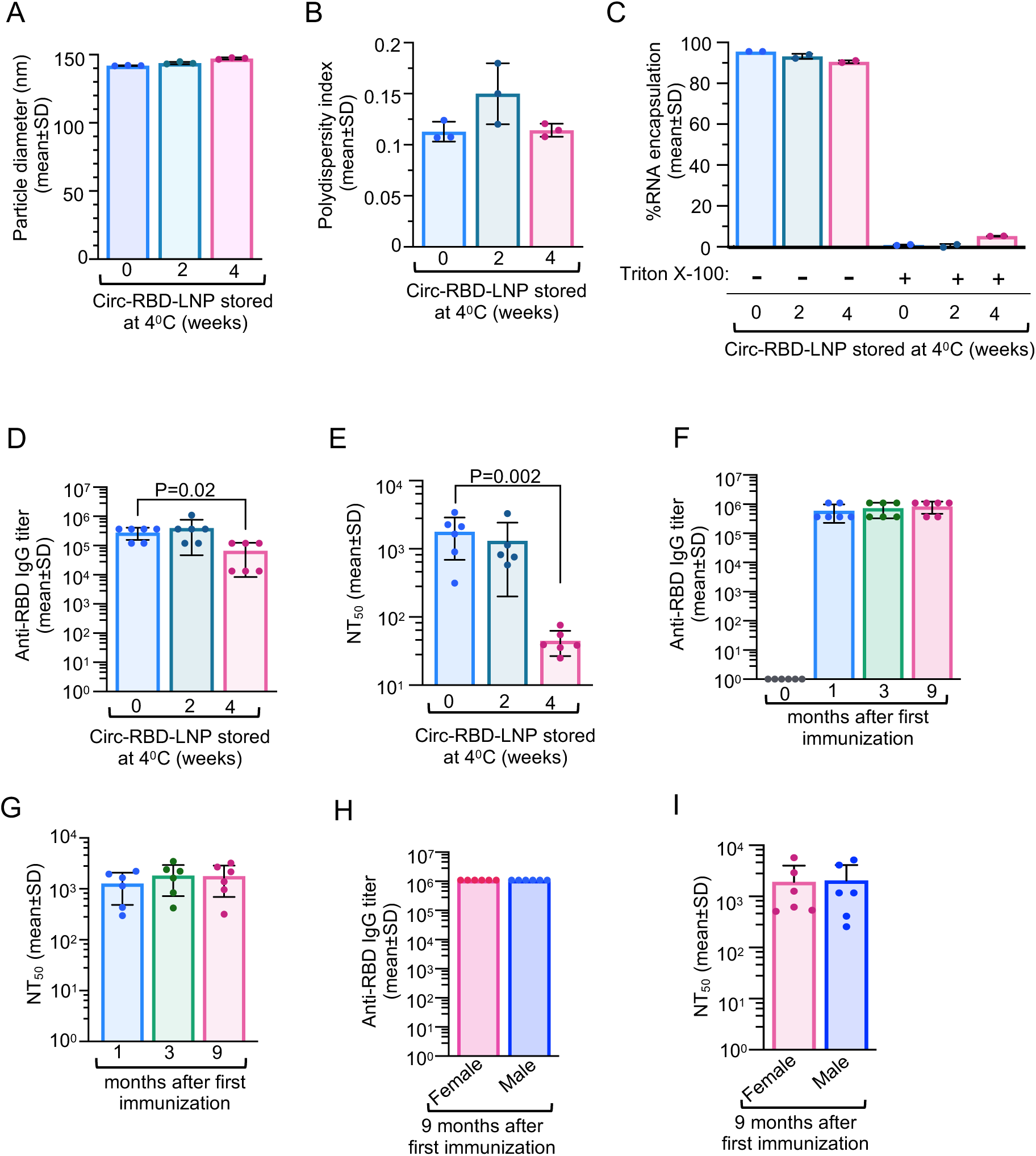
Stability and durability of the immune response induced by the Circ-RBD-LNP vaccine formulation. (A) Average size of the Circ-RBD-LNP stored at 4^0^C for 0, 2 and 4 weeks, evaluated by dynamic light scattering. (B) Average polydispersity index of the Circ-RBD-LNP, stored as indicated in (A). (C) Percentage of RNA encapsulated into the LNP in the Circ-RBD-LNP formulation, stored as indicated in (A). Equal aliquots of the samples were treated with the Triton X-100. (D) Anti-RBD IgG titre in the sera of mice immunized for 4 weeks with the Circ-RBD-LNP, stored as indicated in (A). (E) 50% virus neutralization titre (NT_50_) in aliquots of the sera used in (D), evaluated by PRNT assay. (F) Anti-RBD IgG titre at 0, 1, 3 and 9 months post-immunization in the sera of mice immunized with the Circ-RBD-LNP, evaluated by ELISA. (G) 50% virus neutralization titre (NT_50_) in aliquots of the sera described in (F), evaluated by PRNT assay. (H) Anti-RBD IgG titre at 9 months post-immunization with the Circ-RBD-LNP, in male and female mice, evaluated by ELISA. (I) 50% virus neutralization titre (NT_50_) in aliquots of the sera described in (H), evaluated by PRNT assay. Data are mean±SD of triplicate samples (A-C) and six mice/group (D-I), *p*-values were determined by the unpaired student’s t-test, using the Mann-Whitney test.

Next, durability of the immune response and influence of gender on the immune response elicited by the Circ-RBD was evaluated by immunizing male and female mice with the Circ-RBD vaccine in a two-dose immunization schedule (day 1 and day 14), followed by measurement of the anti-RBD IgG titer and the virus neutralization titer (NT_50_) up to 9 months. Both the anti-RBD IgG titer and the virus neutralization titer (NT_50_) were measured in the mice sera at 1, 3 and 9 months post-first immunization. There was no drop in the anti-RBD IgG titer or the virus neutralization titer (NT_50_) after 9 months of immunization, with reference to the samples collected after one month of the first immunization (Figures 4F and 4G). Both male and female mice induced a similar level of the anti-RBD IgG titer and the virus neutralization titer (NT_50_) (Figures 4H and 4I). Collectively, these results demonstrated that Circ-RBD vaccine is stable in standard refrigeration condition and induced a highly durable neutralizing antibody response in both male and female mice.

### A bivalent SARS-CoV-2 Circ-RBD vaccine induces robust immune response and efficiently neutralizes both the Delta and the Omicron BA.1 variant viruses

Recent reports on bivalent mRNA vaccines and nanoparticle vaccine demonstrate their efficacy in conferring broad spectrum immunity against the SARS-CoV-2 variants of concern (Piccoli et al., 2020; Li et al. 2022; Yuan et al., 2022). Feasibility of using the Circ-RNA vaccine technology for developing a bivalent vaccine was tested by evaluating the immunogenicity and virus neutralization potential of a vaccine formulation containing 1:1 mixture of the Circ-RNAs encoding the RBD of the Delta (Circ-RBD^Del^) and the Omicron BA.1 (Circ-RBD^Omi^) variants of the SARS-CoV-2. Sequence encoding the RBD region of the Delta and the Omicron variants of the SARS-CoV-2 were inserted into the Circ-RNA backbone vector in place of the RBD (Wuhan) sequence so that both RBD^Del^ and RBD^Omi^ are fused in-frame with the CD5 secretory signal and foldon sequence at their N- and C-terminus, respectively. Expression and secretion of the RBD^Del^ and RBD^Omi^ were verified by western blot analysis of the whole cell lysate and the culture medium of the HEK239T cells transfected with the Circ-RBD^Del^-LNP and Circ-RBD^Omi^-LNP formulations. Anti-RBD western blot reveled that both Circ-RBD^Del^ and Circ-RBD^Omi^ were expressed in the HEK239T cells and secreted to the culture medium (Figures 5A, upper panel, 5B, upper and middle panel). Secretion of the Circ-RBD^Omi^ was not as efficient as that of the Delta and the Wuhan variants as the ratio of protein in intracellular to culture medium was much higher in the RBD^Omi^ (Figure 5B, middle panel). GAPDH level was measured in aliquots of the same samples to monitor equal loading (Figures 5A, 5B, lower panel). Mice were injected with vaccine formulations containing 10μg of Circ-RBD^Wuhan^, Circ-RBD^Del^, Circ-RBD^Omi^ or 1:1 mixture of 5μg Circ-RBD^Del^ and 5ug Circ-RBD^Omi^ in a two-dose immunization schedule at two weeks intervals, as described earlier, followed by collection of blood on day 28 and estimation of the anti-RBD IgG titer and the virus neutralization titer. Mice immunized with the vaccines containing the Circ-RBD^Wuhan^, Circ-RBD^Del^ and Circ-RBD^Del+Omi^ produced an anti-RBD IgG titer of approximately 10^6^ against the SARS-CoV-2 Wuhan and Delta variants whereas it was 0.3×10^5^, 0.6×10^5^ and 0.6×10^5^, respectively in case of the Omicron variant (Figures 5C, 5D, 5E). In contrast, the Circ-RBD^Omi^ vaccine induced an average anti-RBD IgG titer of 1.2×10^5^ against the SARS-CoV-2 Wuhan and Delta variants and 1.8×10^5^ against the Omicron variant (Figures 5C, 5D, 5E). The virus neutralization titer (NT_50_) of the sera showed a pattern broadly similar to that of the anti-RBD IgG titer. As observed earlier, NT_50_ of sera from the Circ-RBD^Wuhan^ immunized group against the Wuhan variant was 3.5×10^3^ (Figure 5F). Sera from the Circ-RBD^Del^ and Circ-RBD^Del+Omi^ showed approximately similar NT_50_ as that of the Circ-RBD^Wuhan^ (Figure 5F). However, NT_50_ of sera from the Circ-RBD^Omi^ immunized mice against the Wuhan variant was found to be 0.2×10^3^, which is in line with the data on its anti-RBD IgG titer against the same variant (Figure 5F). NT_50_ of sera from the Circ-RBD^Del^ and the Circ-RBD^Del+Omi^ immunized mice against the SARS-CoV-2 Delta variant was found to be 8.1×10^3^ and 8.8×10^3^, respectively, indicating that the virus neutralization efficacy of the bivalent vaccine was similar to its monovalent counterparts (Figure 5G). Sera from the Circ-RBD^Wuhan^ immunized mice was almost as efficient as sera from the Circ-RBD^Del^ immunized mice in neutralizing the SARS-CoV-2 Delta variant (NT_50_= 1.7×10^3^) (Figure 5G). However, sera from the Circ-RBD^Omi^ immunized mice were significantly less efficient in neutralizing the SARS-CoV-2 Delta variant than that of the Circ-RBD^Wuhan^, Circ-RBD^Del^ and Circ-RBD^Del+Omi^ immunized mice (Figure 5G). Sera from the Circ-RBD^Omi^ and Circ-RBD^Del+Omi^ immunized mice showed an NT_50_ of 8.1×10^3^ and 2.3×10^3^, respectively, against the SARS-CoV-2 Omicron BA.1 variant, indicating that bivalent vaccine was equally effective as its monovalent (Circ-RBD^Omi^) counterpart (Figure 5H). NT_50_ of sera from the Circ-RBD^Del^ and the Circ-RBD^Wuhan^ immunized mice against the Omicron BA.1 variant were found to be 0.3×10^3^ and 0.06×10^3^ respectively, suggesting that they were less efficient in neutralizing the Omicron BA.1 virus (Figure 5H). Collectively, these data demonstrate that the bivalent Circ-RBD vaccine produces potent neutralizing antibodies against the Delta and the Omicron BA.1 variants of the SARS-CoV-2.

**Figure 5.**
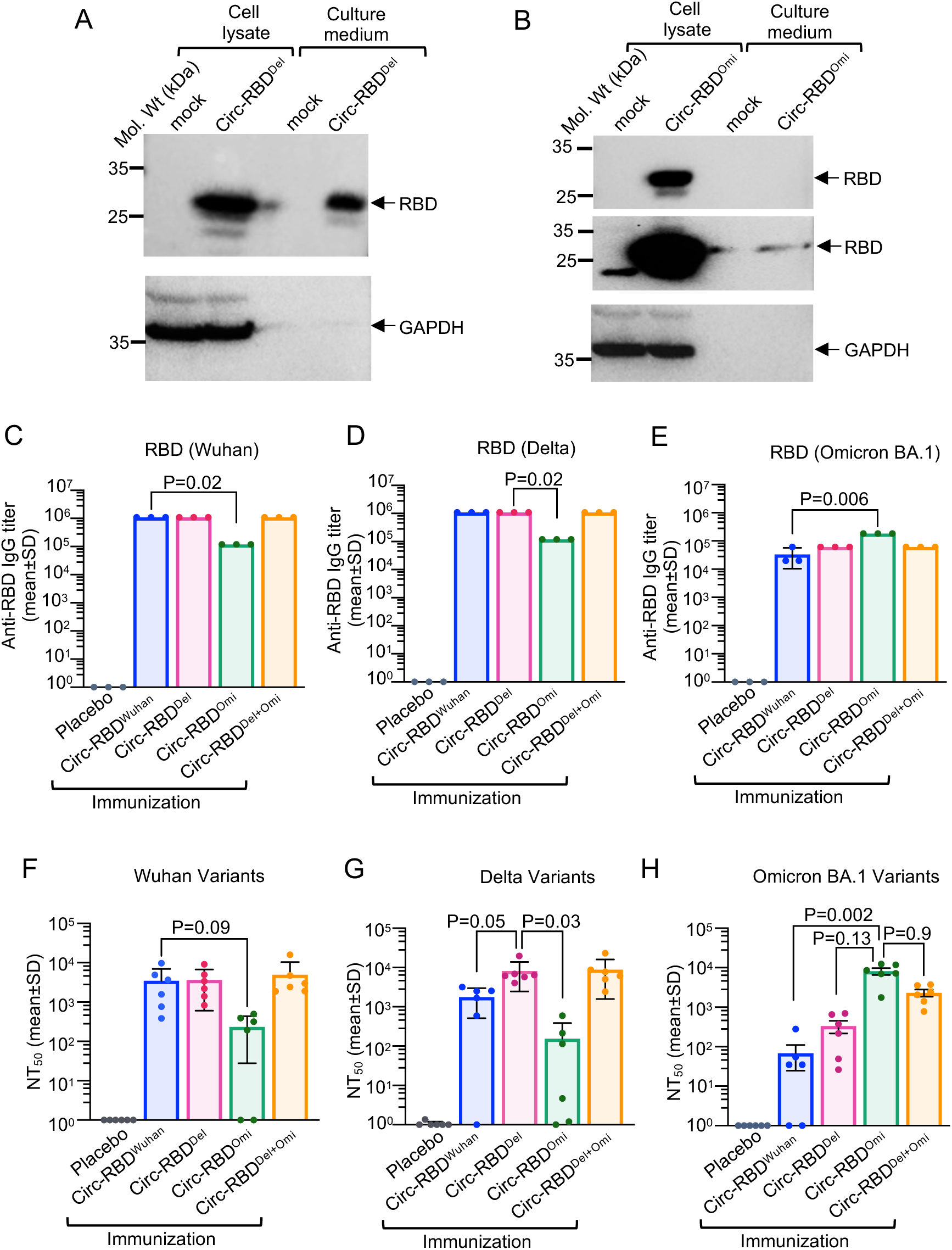
Assessment of immunogenicity and virus neutralization potential of the Circ-RBD^Del^ and Circ-RBD^Omi^ bivalent vaccine formulation. (A) Western blot analysis of the RBD protein in the HEK239T cells transfected with the Circ-RBD-Delta RNA (upper panel). Mock: HEK293T cells treated with LNP. Lower panel: Anti GAPDH western blot of aliquots of the samples used in the upper panel. (B) Western blot analysis of the RBD protein in the HEK239T cells transfected with the Circ-RBD-Omicron-BA.1 RNA (upper panel-lower exposure, middle panel-higher exposure of the same blot). Mock: HEK293T cells treated with LNP. Lower panel: anti GAPDH western blot of aliquots of the samples used in the upper panel. (C) Anti-RBD-wuhan IgG titre at 4 weeks post-immunization in the sera of mice injected with the placebo (PBS), Circ-RBD^Wuhan^, Circ-RBD^Del^, Circ-RBD^omi^ and bivalent Circ-RBD^Del+omi^ vaccine formulations, evaluated by ELISA. (D) Anti-RBD-Delta IgG titre at 4 weeks post-immunization in the sera of mice described in (C), evaluated by ELISA. (E) Anti-RBD-Omicron IgG titre at 4 weeks post-immunization in the sera of mice described in (C), evaluated by ELISA. (F) 50% virus neutralization titre (NT_50_) against the SARS-CoV-2 (Wuhan) at 4 weeks post-immunization in the sera of mice described in (C), evaluated by FRNT assay. (G) 50% virus neutralization titre (NT_50_) against the SARS-CoV-2 (Delta) at 4 weeks post-immunization in the sera of mice described in (C), evaluated by FRNT assay. (H) 50% virus neutralization titre (NT_50_) against the SARS-CoV-2 (Omicron BA.1) at 4 weeks post-immunization in the sera of mice described in (C), evaluated by FRNT assay. In (C)-(E), data are shown as the mean±SD from pooled sera of six mice, assayed in triplicate; in (F)-(H), data are shown as the mean±SD. *p*-values were determined by the non-parametric one-way ANOVA, using the Kruskal-wallis test.

## Discussion

With the success of the COVID-19 mRNA vaccines, immunogenicity and protective efficacy of a number of vaccine candidates based on linear mRNAs, self-amplifying mRNAs and circular mRNAs have been evaluated in multiple independent studies. Given their ability to replicate inside the host, SAM RNAs expresses the antigen for a longer duration with lesser amount of the input RNA. Analogous to the SAM-RNA, Circ-RNAs also offer the advantage of expression of the antigen for a longer duration with lesser amount of the input RNA, owing to their higher stability. On top of that, Circ-RNA vaccines do not produce any additional exogenous protein in the target cells, lack open 5’- and 3’-ends and do not produce double strand RNA (dsRNA) in the target cell, all of which are potential unwanted features in the SAM vaccine. Therefore, Circ-RNA vaccines appear to be more promising than the linear mRNA and SAM vaccines. Comparison between linear mRNA and Circ-RNA vaccines have demonstrated the superiority of the later (Qu et al., 2022). However, there are no report on comparison of the protective efficacy between the Circ-RNA and the SAM vaccines. Here, we compared the efficacy of the Circ-RNA and the SAM vaccines by measuring their immunogenicity, virus neutralization potential and protection efficacy using the SARS-CoV-2 RBD as the vaccine antigen. After confirming that Circ-RBD vaccine induced a better overall response, we further evaluated its thermostability, longevity of the immune response and compatibility for production of bivalent vaccines.

SAM-RNA was produced using the SFV replication machinery. SFV replicon was opted as there are very few reports of SFV infection in human, with very mild symptoms and no instances of mortality. SFV replicon has been used in earlier studies for heterologous protein expression and a recently it has been used to produce a vaccine candidate against the Ebola virus (Liljeström and Garoff, 1991; Krähling et al., 2023). Also note that, multiple outbreaks of VEEV has been reported in North and South American countries with cases of human death. Hence chances of anti-vector immunity is higher in the case of VEEV (Weaver et al., 1996). The Anabaena permuted intron-exon sequence was used to produce the Circ-RNA and CVB3 IRES-mediated translation was used to produce the secretory signal and foldon fused-RBD protein in the target cells. Circ-RNA vaccine production using the above vector has been reported and well characterized (Qu et al., 2022). Only the RBD region was used instead of the entire Spike protein of the SARS-CoV-2 as it contains the majority of neutralizing epitopes, avoids induction of antibody-dependent enhancement (ADE) and permits insertion of multiple elements due to its smaller size. The trimeric structure of RBD was maintained using the foldon-motif to improve its immunogenicity and virus neutralization potential, as reported earlier (Routhu et al., 2021).

In order to compare the SAM-RBD vaccine with the Circ-RBD vaccine, it was important that both vaccine antigens were expressed at comparable levels and did not induce the host innate immune response. Western blot analysis using anti-RBD antibody and RT-qPCR analysis of the ISGs confirmed that both vaccine candidates expressed similar levels of the vaccine antigen and evaded the host innate immune pathway RNA sensors, respectively.

Although both SAM-RBD and Circ-RBD induced a similar humoral response, CD4^+^ and CD8^+^ memory T cell response was significantly increased in the Circ-RBD immunized mice. There was a trend of increased production of IFNγ and TNFα in CD4+ and CD8+ memory T cells in SAM-RBD immunized mice, however, it was not statistically significant, indicating that Circ-RBD vaccine induced a more potent cell-mediated immune response. SFV replicon system has been used to develop a dual antigen vaccine candidate against the Ebola virus by co-expressing the viral glycoprotein and nucleoprotein where it was observed that high intradermal dose of the Ebola virus glycoprotein induced IFNγ in CD4+ T cells and high intradermal dose of nucleoprotein induced very low level of IFNγ in CD8+ T cells (Krähling et al., 2023). In the case of VEEV-replicon based SARS-CoV-2-RBD and nucleocapsid (N)-expressing dual antigen vaccine immunized outbred mice, IFNγ level was moderately increased in the CD4+ and CD8+ T cells stimulated with either the S- or N-peptide pool (McCafferty et al., 2022). Other studies on evaluation of a SAM vaccine candidate in mice based on the SARS-CoV-2-RBD or the Spike protein expressed from an alpha virus (VEEV/ VEEV+Sindbis) replicon reported a significant T-cell response in splenocytes of the immunized mice, upon stimulation with the S-peptide pool (McKay et al., 2020; Erasmus et al., 2020; de Alwis et al., 2021; Maruggi et al., 2022). However, SAM-Spike immunized pigtail macaques did not induce high level of IFNγ production in CD4+ and CD8+ T cells upon stimulation of splenocyte with S-peptide pool (Erasmus et al., 2020). On the other hand, the Circ-RBD vaccine of the SARS-CoV-2 induced similar level of T cell response as that of the corresponding linear mRNA vaccine and a higher neutralizing antibody response than that of the linear mRNA (Qu et al., 2022). Given that different assays were conducted in different laboratories to measure the T cell response, it is possible that lower IFNγ and TNFα production in CD4+ and CD8+ T cells (splenocytes stimulated with RBD peptide pool) may be attributed to the overall lesser sensitivity of our assay protocol. However, the relative difference in the level of IFNγ and TNFα between SAM-RBD and Circ-RBD immunized sample would remain constant as the samples were processed in parallel. Therefore, Circ-RBD vaccine appears to be a better alternative to a SAM-RBD vaccine. Whether a VEEV replicon-based SAM vaccine shows a similar profile as the SFV replicon used in our study, remains to be tested. A recent study also showed potent adjuvant function of the circular RNA, irrespective of the antigen sequence and robust CD8+ T-cell response induced by a circular RNA vaccine encoding the nucleoprotein of the influenza virus, further supporting practical utility of the circular RNA vaccine technology (Amaya et al., 2023).

Functional efficacy of the Circ-RBD vaccine was characterized by evaluating its thermostability, durability of the neutralizing antibody response, ability to protect the mice upon SARS-CoV-2 challenge and compatibility as a bivalent vaccine formulation. Circ-RBD immunization induced peak anti-RBD IgG titer at 4 weeks, which protected the mice from COVID-19 like disease pathology when SARS-CoV-2 infection was initiated in the lower respiratory tract (severe disease, often leading to lethality). Nine month follow up study of the anti-RBD IgG titer and the virus neutralization titer (NT_50_) did not show any decline in both. Moreover, neutralizing antibody titer was similar in both male and female mice. With regard to stability of the Circ-RBD vaccine in normal refrigeration conditions, there was no change in the characteristic of the Circ-RBD-LNP formulation after 4 weeks of storage, however neutralizing antibody titer was stable for up to 2 weeks of storage. Finally, we evaluated the functional efficacy of a bivalent Circ-RBD vaccine. Failure of monovalent vaccines in preventing COVID-19 for a longer period is ascribed to emergence of newer variants of the SARS-CoV-2 containing escape mutations in the RBD region. Bivalent vaccines based on linear mRNA platform or nanoparticle platform expressing the spike protein or RBD region of the spike protein of the SARS-CoV-2-Delta and Omicron variants have shown cross-protection against SARS-CoV-2 variants (Corleis et al., 2023; Li et al., 2022; Yuan et al. 2022). On similar line, our data on the bivalent Circ-RBD vaccine expressing the RBD of Delta and Omicron variants demonstrate cross-protection against the Wuhan, Delta and Omicron variants.

In summary, the current study compared the immunogenicity and virus neutralization efficacy of the SAM and Circ-RNA vaccine candidates encoding the SARS-CoV-2-RBD antigen, which revealed similar humoral immune response and virus neutralization potential of both vaccines and a superior cell-mediated immune response induced by the later. Our data also showed reasonable stability of the Circ-RNA vaccine at four degrees centigrade, high durability of the neutralizing antibody response induced by it and high virus neutralization efficacy of the bivalent Circ-RNA vaccine, all of which support the use of Circ-RNA technology in vaccine development. An earlier study showed the superiority of the SARS-CoV-2 Circ-RNA vaccine over the corresponding linear mRNA vaccine. Data obtained from this study further supports the use of the Circ-RNA technology for vaccine development.

## Materials and Methods

### Plasmids and reagents

Plasmid encoding the noninfectious Semliki Forest virus replicon (pSFV3) was a gift from Henrik Garoff (Addgene plasmid # 92072; http://n2t.net/addgene:92072; RRID:Addgene_92072) (Liljeström and Garoff, 1991). 988 to 1578 nucleotides encoding the RBD region (330 to 526 amino acids) of spike protein of the SARS-CoV-2 Wuhan isolate (hCoV-19/USA-WA1/2020) fused in-frame with the CD5 secretory sequence at the N-terminal and the Bacteriophage T4 fibritin trimerization motif sequence at the C-terminus was synthesized and inserted into the pSFV3 vector at the BamH1 restriction site. The resulting plasmid was named pSFV3-RBD. Transcription of SFV-RBD hybrid RNA in the pSFV3-RBD vector was under control of the SP6 promoter. A circular RNA expression cassette containing the sequence of the T7 promoter, Anabaena Group 1 intron 3’ sequence, CVB3 IRES sequence, CD5 secretory sequence, RBD region (988 to 1578 nucleotides) of spike protein of the SARS-CoV-2 Wuhan isolate (hCoV-19/USA-WA1/2020), bacteriophage T4 fibritin trimerization motif and anabaena group 1 intron 5’ sequence was chemically synthesised in the given order and cloned into the pUC57 vector. The resulting plasmid was named pUC57 cRNA-RBD^Wuhan^. RBD corresponding to the SARS-CoV-2-Delta (hCoV-19/USA/MD-HP05285/2021) and Omicron (hCov-19/USA/MD-HP2087/2021) variants were cloned similarly and the resulting plasmids were named pUC57 cRNA-RBD^Del^ and pUC57 cRNA-RBD^Omi^, respectively. The goat anti-Mouse IgG-HRP (Catalog No. 1030-05) was from Southern Biotech (Alabama, USA). Megascript SP6 transcription kit (Catalog no. AM1330), MegaScript T7 transcription kit (Catalog no. AM1334) lipofectamine 3000 (Catalog No. L3000075), anti-SARS-CoV-2 Spike S1 (Catalog no. PA5-81795), goat anti-Rabbit IgG-HRP (Catalog No. 31460), Anti-mouse IgG (H+L) Alexa Flour 488 (Catalog no. A28175) and Ribogreen (Catalog no. R11490) were from Thermo Fisher Scientific (Massachusetts, USA). Scriptcap Cap 1 capping system (Catalog no. sccs1710) was from CellScript (Wisconsin, USA). RNAse R (Catalog no. ab286929) was from Abcam (Cambridge, United Kingdom). RNeasy Kit was from Qiagen ((Catalog no. 74104, Hilden, Germany). SM102 (Catalog no. 33474) was from Cayman Chemical company (Michigan, USA). DSPC (Catalog no. 850365P), Cholesterol (Catalog no. 700000P), and DMG-PEG 2000 (catalog no. 880151P), was from Sigma (Missouri, USA). Amicon ultra centrifugal filter (Catalog no. UFC703008) was from Merck Millipore (Darmstadt, Germany). anti-GAPDH antibody (Catalog no. SC-25778) was from Santa Cruz Biotechnology (Texas, USA). TRI-reagent (Catalog no. TR118) was from MRC (Massachusetts, USA). Firescript cDNA synthesis kit (Catalog no. 06-15-00050) and HOT FIREPol SolisGreen qPCR Mix (Catalog no.08-47-0000S) from Solis Biodyne (Estonia). Ad5-hACE (catalog no. VVC-McCray-7723) was obtained from the Viral Vector Core (University of Iowa, Iowa, USA). Vetbond (catalog no. 1469Sb) was from 3M (Minnesota, USA). Mouse monoclonal antibody isotyping reagent (Cat no. ISO2) was from Sigma Aldrich (Missouri, USA). SARS-CoV-2 spike antibody (catalog no. GTX632604) was from GeneTex (California, USA). SARS-CoV-2 RBD peptide pool (Catalog no: PM-WCPV-S_RBD-1) was from JPT innovative peptide solution (Berlin, Germany). GolgiStop (Catalog no. 554724) was from BD bioscience (New Jersey, USA). FC (Catalog no.14-9161-73), IC fixation buffer (Catalog no. FB001), permeabilizing buffer (Catalog no. 00-8333-56), anti-mouse CD3 (Catalog no.100204), anti-mouse CD4 (Catalog no.100408), anti-mouse CD8 (Catalog no.100714), anti-mouse CD44 (Catalog no.103030), anti-mouse IFNγ (Catalog no. 505810), TNFα (Catalog no. 506328), and IL-4 (Catalog no. 504126) are from Thermo Fisher Scientific (Massachusetts, USA).

### *In vitro* transcription, capping, circularization, RNase R treatment and purification

pSFV3-RBD, pUC57 cRNA-RBD^Wuhan^, pUC57 cRNA-RBD^Del^ and pUC57 cRNA-RBD^Omi^plasmids were linearized and purified DNA was used for the *in vitro* transcription using the Megascript SP6 or T7 transcription kit, following the manufacturer’s instruction. SAM-RNA was capped using the Scriptcap Cap 1 capping system, following the manufacturer’s instructions. In the case of Circ-RNA, *in vitro* transcribed RNA was incubated at 55^0^C for 15 min in the circularization buffer (10mM MgCl_2_, 50mM Tris-HCl, 100mM NaCl, 1mM DTT, pH 7.9; New England Biolabs, USA) and 2μM rGTP to circularize the RNA, as described earlier (Wesselhoeft et al., 2018b). Circularized RNA was incubated at 37^0^C for 7 min with 0.1units RNase R/μg RNA, followed by addition of 0.1 units RNase R/μg RNA and incubation for another 7 min. RNA was purified using the RNeasy kit, following the manufacturer’s protocol (Qiagen, Germany).

### Preparation and characterization of the RNA-LNP vaccine formulation

SM102, DSPC, Cholesterol and DMG-PEG 2000 were mixed at a molar ratio of 50:10:38.5:1.5 in absolute ethanol at 1/5 of the final LNP amount. An aqueous solution of RNA was prepared by diluting it in 50mM acetate buffer (1.2mM sodium acetate, 4mM acetic acid, pH 4.0) at 4/5 of the final LNP volume. The two solutions were mixed using a microfluidics system (ElveFlow Elvesys, France) at a flow rate ratio of 1:5. The resulting RNA-LNP complex was dialyzed with phosphate buffered saline (137 mM NaCl, 2.7 mM KCl, 10 mM Na_2_HPO_4_, 1.8 mM KH_2_PO_4_, pH 6.8) and concentrated using a 30kDa MWCO Amicon ultra centrifugal filter to a volume of 0.1μg/μl RNA concentration. Further, the RNA-LNP vaccine formulation was passed through a 0.2μm filter and aliquoted into glass vials and stored at 4^0^C. Size and polydispersity (PDI) of the RNA-LNP vaccine formulation was evaluated by dynamic light scattering (ZetaSizer lab, Malvern panalytical, United Kingdom). For transmission electron microscopy (TEM), RNA-LNP vaccine formulation was diluted ten times in H_2_O, followed by staining with 2% uranyl acetate and image acquisition using JEM-1400 Flash TEM equipped with tungsten filament and sCMOS camera (JEOL Ltd, Japan). RNA encapsulation into LNP was measured by fluorescent-based RNA quantification system using Ribogreen, following manufacturer’s instruction. The encapsulated RNA was released from the LNP by treating the RNA-LNP vaccine formulation with 2% Triton X-100, followed by measurement of RNA quantity by Ribogreen assay and evaluation of RNA quality by agarose gel electrophoresis.

### Mammalian cell culture, transfection, western blot, RT-qPCR, SARS-CoV-2 propagation

HEK293T cells were maintained in Dulbecco’s modified Eagle medium (DMEM) supplemented with 10% FBS (fetal bovine serum) and Penicillin, Streptomycin during maintenance. Cells were seeded on 6 well plates (8×10^5^ cell/well) and transfected with 6μg of RNA-LNP complex. Only LNP was added in the mock. For western blot analysis under reducing condition, samples (cells or culture medium) were incubated at 95^0^C for 5 minutes in Laemlli buffer (2% SDS, 13% glycerol, 2.5% β-mercaptoethanol, 0.005% bromo phenol blue and 63 mM Tris HCl, pH 6.8). For non-reducing condition, β-merceptoethanol was omitted from the Laemlli buffer. Total protein in the samples was quantified by Bicinchoninic acid assay and equal amount of protein were resolved in SDS-PAGE, followed by transfer to Polyvinylidene fluoride (PVDF) membrane. The membrane was blocked with 5% skimmed milk, incubated with anti-Spike S1 or anti-GAPDH antibody at 1:5000 dilution. Anti-Rabbit secondary antibodies (Santa Cruz Biotechnology) were used at 1:5000 dilution. Protein bands were developed by enhanced chemiluminescence reagent, using a commercially available kit (BioRad, USA) and visualized in a gel Documentation system (Chemidoc MP, Bio Rad, USA).

For RT-qPCR analysis of genes involved in the antiviral response pathway, HEK293T cells were transfected with 1μg of SAM-RBD or Circ-RBD RNA or Poly I:C was using lipofectamine 3000, following manufacturer’s instruction (Thermo Fisher Scientific, Massachusetts, USA). 24 hours post-transfection, total RNA was isolated using the TRI-reagent (MRC, Massachusetts, USA), followed by reverse transcription (RT) using the Firescript cDNA synthesis kit (Solis Biodyne, Estonia). The relative transcript levels of different genes were determined by SYBR green based RT-qPCR, as described earlier (Verma et al., 2021). Values obtained for GAPDH RNA was used as an internal control for normalization of the data. Primer sequences are as follows: hGAPDH FP: 5’ GAGTCAACGGATTTGGTCGT 3’, hGAPDH RP: 5’ TTGATTTTGGAGGGATCTCG 3’, hIFNβ FP: 5’ ACGCCGCATTGACCATCTAT 3’, hIFNβ RP: 5’ TTGGCCTTCAGGTAATGCAGA 3’, hRIG-I FP: 5’ TGTGGGCAATGTCATCAAAA 3’, hRIG-I RP: 5’ GAAGCACTTGCTACCTCTTGC 3’, hOASL FP; 5’ AGGGTACAGATGGGACATCG 3’, hOASL RP: 5’ AAGGGTTCACGATGAGGTTG 3’, hPKR FP: 5’ TCTTCATGTATGTGACACTGC 3’, hPKR RP: 5’ CACACAGTCAAGGTCCTT 3’, hOAS1 FP: 5’ GCTCCTACCCTGTGTGTGTGT 3’, hOAS1 RP: 5’ TGGTGAGAGTACTGAGGAAGA 3’.

For RT-qPCR analysis in the mouse lung tissue, total RNA was isolated from 10mg lung tissue using TRI reagent, following manufacturer’s instruction. cDNA synthesis and RT-qPCR was done as mentioned above. HPRT RNA level was quantified for normalization of data. Primer sequences are as follows: SCoV2 FP: 5’ TGGACCCCAAAATCAGCGAA 3’, SCoV2 RP: 5’ TCGTCTGGTAGCTCTTCGGT 3’, mHPRT FP: 5’ GTTGGATACAGGCCAGACTTTGTTG 3’, mHPRT RP: 5’ GATTCAACTTGCGCTCATCTTAGGC 3’, mIL-6 FP: 5’ GAGGATACCACTCCCAACAGACC 3’, mIL-6 RP: 5’ AAGTGCATCATCGTTGTTCATACA 3’, mIL-1β FP: 5’ GCCCATCCTCTGTGACTCAT 3’, mIL-1β RP: 5’ AGGCCACAGGTATTTTGTCG 3’, mTNFα FP: 5’ CATCTTCTCAAAATTCGAGTGACAA 3’, mTNFα RP: 5’ TGGGAGTAGACAAGGTACAACCC 3’, mIFNγ FP: 5’ AACGCTACACACTGCATCTTGG 3’, mIFNγ RP: 5’ GACTTCAAAGAGTCTGAGG 3’.

SARS-CoV-2 Wuhan isolate (NR-52281, SARS-related coronavirus-2 isolate ISA-WA1/2020) and Omicron Variants (B.1.1.529, NR56461, SARS-related coronavirus-2 isolate hCoV-19/USA/MD-HP20874/2021) were obtained from BEI Resources, amplified in Vero E6 cells in the BSL3 facility of THSTI, India, titrated and stored frozen in aliquots. SARS-CoV-2-Delta (SARS-CoV-2/human/IND/THSTI_287/2021) was isolated from clinical sample at THSTI and has been reported (GenBank: MZ356566.1). SARS-CoV-2 infection was done as described^43^.

### Mice handling, immunization, SARS-CoV-2 challenge study

Mice were housed in individually ventilated cages at the experimental animal facility of THSTI. For SARS-CoV-2 infection experiments, animals were housed in the animal biosafety level-3 (ABSL-3) laboratory of THSTI. Institutional animal ethics committee and institutional biosafety committee approval was obtained for the experiments. Male and Female Balb/c mice of 6 to 8 weeks old were intradermally injected on the back of the animal. In the case of intramuscular injection, mice were injected in the hind leg muscle. All injections were in 100μl volume. Placebo animal received 100μl PBS. Blood was collected at the indicated time points through retro-orbital sinus using a sterile hematocrit capillary tube. Wherever indicated, mice were euthanized by anesthetic overdose (Ketamine 300 mg/kg and xylazine 30 mg/kg) and blood, spleen and lungs were collected. Serum was isolated from the blood by centrifugation at 10,000g for 10 min at 4°C. Lung tissue was cut into parts and used for RNA, protein or histopathology analysis.

SARS-CoV-2 challenge experiments were conducted in inbred Balb/c mice. Six to eight-week-old mice were immunized with 10μg of Circ-RBD twice at two-weeks interval. Blood was collected through retro-orbital sinus and anti-RBD IgG was measured. Replication deficient adenovirus expressing human ACE2 (Ad5CMV-hACE2, denoted as hACE2) was transduced through intranasal or intratracheal route (Hassan et al., 2020; Gu et al., 2021). For Intranasal transduction, mice were lightly anaesthetized with isoflurane and 2×10^8^ pfu of hACE2 was administered, as described (Hassan et al., 2020). Intratracheal transduction of hACE2 was performed as described (Gu et al., 2021). Mice were anaesthetised with ketamine xylazine injection (Ketamine 100 mg/kg and xylazine 10 mg/kg), hair was shaved from the neck region and a 5mm incision was made in the skin to expose the trachea. 2×10^8^ pfu of hACE2 was administered in the trachea using a 24-gauge I.V. cannula. The wound opening was glued with tissue adhesive (Vetbond, 3M Corp, USA). Animals did not show any sign of illness or inflammation at the site of glue. Mice were infected with 10^5^ pfu SARS-CoV-2 intranasally after 5 days of hACE2 administration. Weight of the animals were measured before infection and every day after the infection. In the intranasal transduction model, mice were sacrificed 5 days post-SARS-CoV-2 infection and lung and BAL fluid were harvested. In the case of intratracheal transduction model, weight was measured every day after infection and survival was checked till 21 days of post-infection. When an animal showed critical illness symptoms after SARS-CoV-2 infection, it was euthanized and samples were collected. An animal from the immunized group was euthanized for comparison when mice from the placebo group were euthanized.

### ELISA, Flow cytometry, quantification of viral load, virus Neutralization assay

The Enzyme-Linked Immunosorbent Assay (ELISA) was performed as described, with below mentioned modifications (Mehdi et al., 2021). The receptor-binding domains (RBDs) of the three SARS-CoV-2 variants-Wuhan, Delta, and Omicron, were expressed in-house in the Expi293 cells (Thermo Fisher Scientific, Massachusetts, USA) and used in the ELISA. The assay samples (mice sera) were diluted in a 3-fold series from 1:500 to 1:1,093,500. Horse radish peroxidase (HRP)-labelled Goat anti-Mouse IgG (H+L) (Jackson ImmunoResearch, USA) was used at a dilution of 1:5000. The endpoint anti-RBD IgG titer (denoted as anti-RBD IgG titer in the text and figures) refers to the dilution factor at which the absorbance (obtained in the ELISA) of the test samples exceeded three times that of the background (secondary Ab alone). Anti-RBD IgM and IgG isotype (IgG1, IgG2a and IgG2b) ELISA was performed using a commercially available kit, following the manufacturer protocol (Sigma Aldrich) at a serum dilution of 1:500.

For FACS study, splenocytes were prepared as described and cultured in RPMI-10 supplemented with 10%fetal bovine serum and 1% penicillin-streptomycin antibiotic (Agarwal et al., 2022). After 3 hours, RBD peptide pool (2 μg/ml each peptide) and GolgiStop (BD) was added to the culture medium for 6 hours. Cells were blocked with FC at room temperature for 15 min, stained with anti-CD3, anti-CD4, anti-CD8 and anti-CD44 antibodies followed by fixation and permeabilization. Intracellular staining of IFNγ, TNFα and IL-4 was performed using respective antibodies. Cells were sorted using the BD FACS-aria (BD Biosciences) and data was analyzed using the FlowJo software (FlowJo LLC, Oregon, USA).

For quantification of SARS-CoV-2 level in the lungs, 5mg of lung tissue was homogenised in 1ml serum free DMEM and sterilized using 0.2μm filter. The homogenate was serially diluted 10-fold and added to 2×10^5^ VeroE6 cells/well in 12 well plate for 1 hour. Infection medium was removed, 1% carboxymethyl cellulose (CMC) was added and plates were incubated for 48 hours at 37^0^C, with 5% CO_2_ supplementation. Cells were fixed in 3.7% formaldehyde for overnight, plaques stained with 0.05% crystal violet and number of plaques in each well was counted.

50% virus neutralization efficiency (NT_50_) of the sera from mice immunized with the SAM-RBD^Wuhan^ and Circ-RBD^Wuhan^ vaccine formulations (shown in Figs. 2D, 3E, 3F, 3H, 3J, S1B and S1D) were measured by the Plaque reduction neutralization test (PRNT), as described (Bewley et al., 2021). Briefly, mouse sera were heated at 55^0^C for 30 min, 2-fold serially diluted starting from 1:10 in serum-free DMEM, mixed with an equal volume of virus [virus stock was appropriately diluted so that each well produces approximately 50 plaque forming units (pfu)] and incubated at 37^0^C for 1 hour. 2×10^5^ VeroE6 cells/well in 12 well plates were incubated with the serum-virus mix for 1 hour. Infection medium was removed and cells were overlaid with 1% CMC in 2% FBS supplemented DMEM and incubated at 37^0^C with 5%CO_2_ supplementation for 48 hours. Cells were fixed in 3.7% formaldehyde for overnight and stained with 0.2% crystal violet. The number of plaques were counted and the serum dilution leading to 50% neutralization of the virus (NT_50_) was calculated using the GraphPad Prism 9.0 (Massachusetts, USA). NT_50_ of the Wuhan, Delta and Omicron variants (shown in Figures 6F-6H) were measured by the Focus reduction neutralization test (FRNT). Heat inactivated serum sample were 2-fold serially diluted, starting from 1:20 and mixed with an equal volume of the virus [virus stock was appropriately diluted so that each well produces approximately 100 focus forming units (ffu)] and incubated at 37^0^C for 1 hour. 2×10^4^ VeroE6 cells/well in 96 well plates were incubated with the serum-virus mix for 1 hour. Infection medium was removed and cells were overlaid with 0.5% CMC in 2% FBS supplemented DMEM and incubated at 37^0^C with 5%CO_2_ supplementation for 24 hours. Cells were fixed in formaldehyde and permeabilized with 0.2% Triton X-100. Next, the cells were stained with anti-Spike antibody, following manufacturer’s protocol followed by staining with alexa fluor-488-conjugated anti-mouse IgG antibody. The number of foci was counted using an ELISPOT reader (iSPOT, AID GmBh, Strassberg, Germany) and the serum dilution leading to 50% neutralization of the virus was calculated using GraphPad Prism 9.0.

### Histopathology

Histopathology analysis of the lung tissue samples were performed by the histology service of the experimental animal facility of THSTI, following established protocol. Briefly, lung tissues were kept in 10% neutral buffered formalin solution immediately after collection. After ensuring proper fixation, samples were subjected to processing and paraffin embedded blocks were prepared. 4μm thick sections were cut using a semi-automatic microtome (Histocore Multicut-Leica Biosystem, Germany) and mounted on glass slides. The sections were stained with haematoxylin and eosin, following standard operating procedures of the facility. Slides were analysed by a qualified histopathologist, following blind evaluation method. Semi-quantitative scoring system was adopted to score the pathological conditions, 0 indicating absence of a pathological condition while 5 indicating its highest severity. Pathological conditions such as; bronchiolitis, vasculitis, interstitial pneumonia, alveolitis and haemorrhage were analysed and scored. Images were visualized and captured using a Bright field upright microscope (CX43RF-Olympus Life science Solutions, Japan) equipped with a colour camera (DP28-Olympus Life science Solutions, Japan) and imaging software (cellSens Entry-Olympus Life science Solutions, Japan) at 200X magnification.

### Statistical analysis

All cell line data are derived from 3 independent experiments and represented as mean±SD of 3 independent experiments. For mouse study, 6 or more animals were used in each group, as indicated in the figures. Average values are shown in the bar and individual values are shown as dots in each bar in the graphs. Two-tailed unpaired t*-*test (Mann-Whitney) and One-way ANOVA (Kruskal-Wallis test) followed by Dunn’s multiple comparisons was used for comparison between groups, as indicated in the figures. Logistic regression was used to determine the NT_50_ titre of the mice sera. All statistical analysis was performed using the GraphPad Prism 9. A *p*-value <0.05 with 95% confidence interval was considered statistically significant.

## Acknowledgement

This study was partly supported by a BIRAC (Biotechnology Industry Research Assistance Council, Department of Biotechnology, Government of India) grant to MS (BT/COVID0081/01/20) and THSTI core grant to MS. ONS and UB are supported by senior research fellowships from the Council of scientific and industrial research and the department of Biotechnology, Government of India, respectively. We are thankful to the experimental animal facility and biosafety level three (BSL3) facility of THSTI for helping with the experiments. The following reagent was deposited by the Centre for Disease Control and Prevention (USA) and obtained through BEI Resources, NIAID, NIH (USA): SARS-Related Coronavirus 2, isolate hCoV-19/USA-WA1/2020, NR-52281 and SARS-Related Coronavirus 2, isolate hCoV-19/USA/MD-HP20874/2021 (Lineage B.1.1.529; Omicron Variant), NR-56461, contributed by Andrew S. Pekosz. We are thankful to Henrik Garoff for the pSFV3 plasmid.

## Author contribution

Conceptualization: MS; Investigation: ONS, UB, GJ, TRA, KC, SN, PS, SC, FM; Data analysis: ONS, GJ, TRA, BNP, SM, GB, CTRK, PG, MS; Resources: TS, SM, DN, GB, CTRK, PG, MS; manuscript writing: ONS, MS, GB, SM, CTRK. All authors have reviewed, edited and agreed to the final version of the manuscript.

---

**Figure S1.**
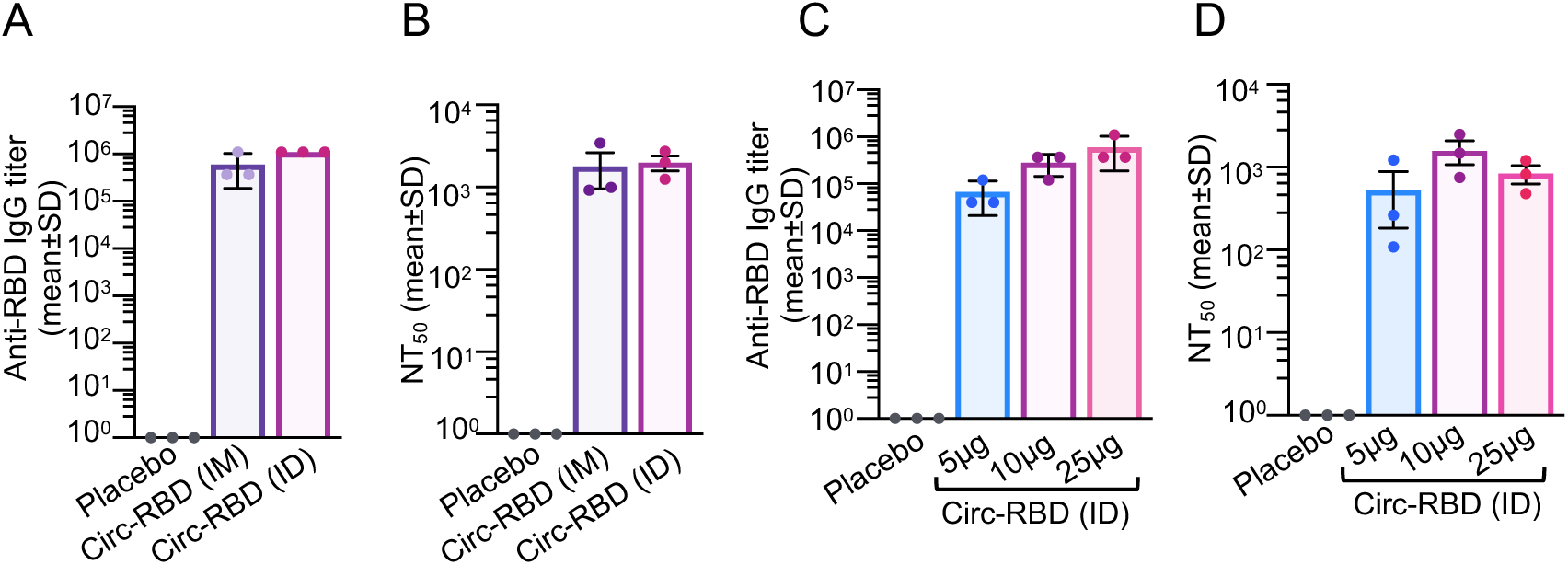
Selection of immunization route and dose of RNA-LNP formulation that elicits significant immune response in mice. (A) Anti-RBD IgG titre at 4 weeks post-immunization in the sera of mice injected with the placebo (PBS) and the Circ-RBD-LNP via intramuscular (IM) or intradermal (ID) route, evaluated by ELISA. (B) 50% virus neutralization titre (NT_50_) at 4 weeks post-immunization in the sera of mice described in (B), evaluated by PRNT assay. (C) Anti-RBD IgG titre at 4 weeks post-immunization in the sera of mice intradermally injected with the placebo (PBS) or 5, 10, 25μg of Circ-RBD-LNP, evaluated by ELISA. (D) 50% virus neutralization titre (NT_50_) at 4 weeks post-immunization in the sera of mice described in (C), evaluated by PRNT assay. Data are shown as the mean±SD of pooled sera from five animals, assayed in triplicate; *p*-values were determined by the unpaired student’s t-test, using the Mann-Whitney test.

**Figure S2.**
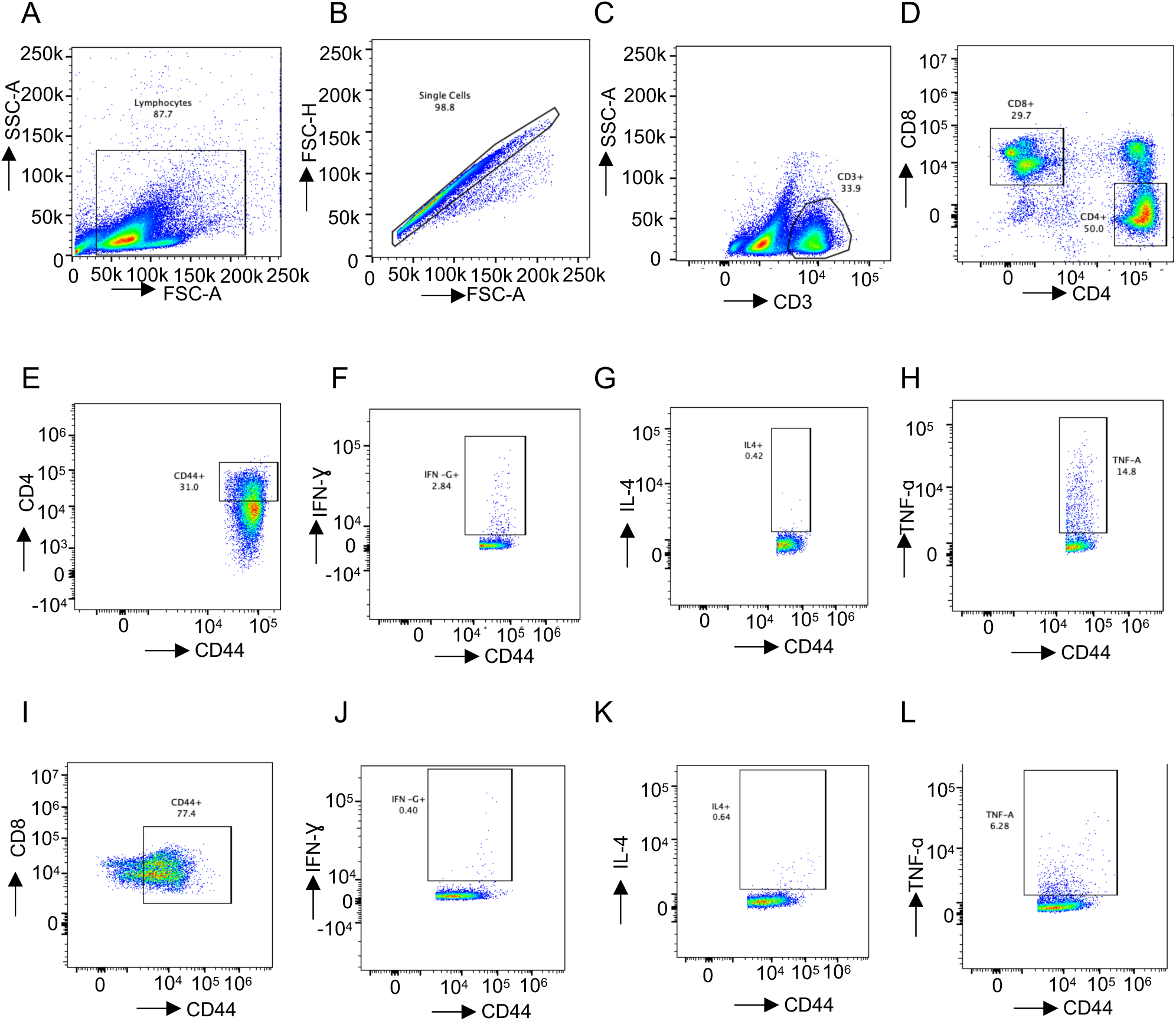
Analysis of intracellular IFNγ, TNFα and IL-4 levels in the RBD peptide pool-stimulated CD4, CD8 positive T-cells derived from the splenocytes of Circ-RBD immunized mice. (A)-(D) Representative FACS plot showing the gating strategy of lymphocytes (A), single cells (B), CD3^+^ (C) and CD4^+^CD8^+^(D) T-cells. (E)-(H) Representative FACS plot showing the gating strategy of CD4^+^ memory T-cells (E) and quantification of IFNγ (F), IL-4 (G) and TNFα (H) in CD4^+^ memory T-cells. (I)-(L) Representative FACS plot showing the gating strategy of CD8^+^ memory T-cell (J) and quantification of IFNγ (K), IL-4 (K) and TNFα (L) in CD8^+^ memory T-cells.

**Figure S3.**
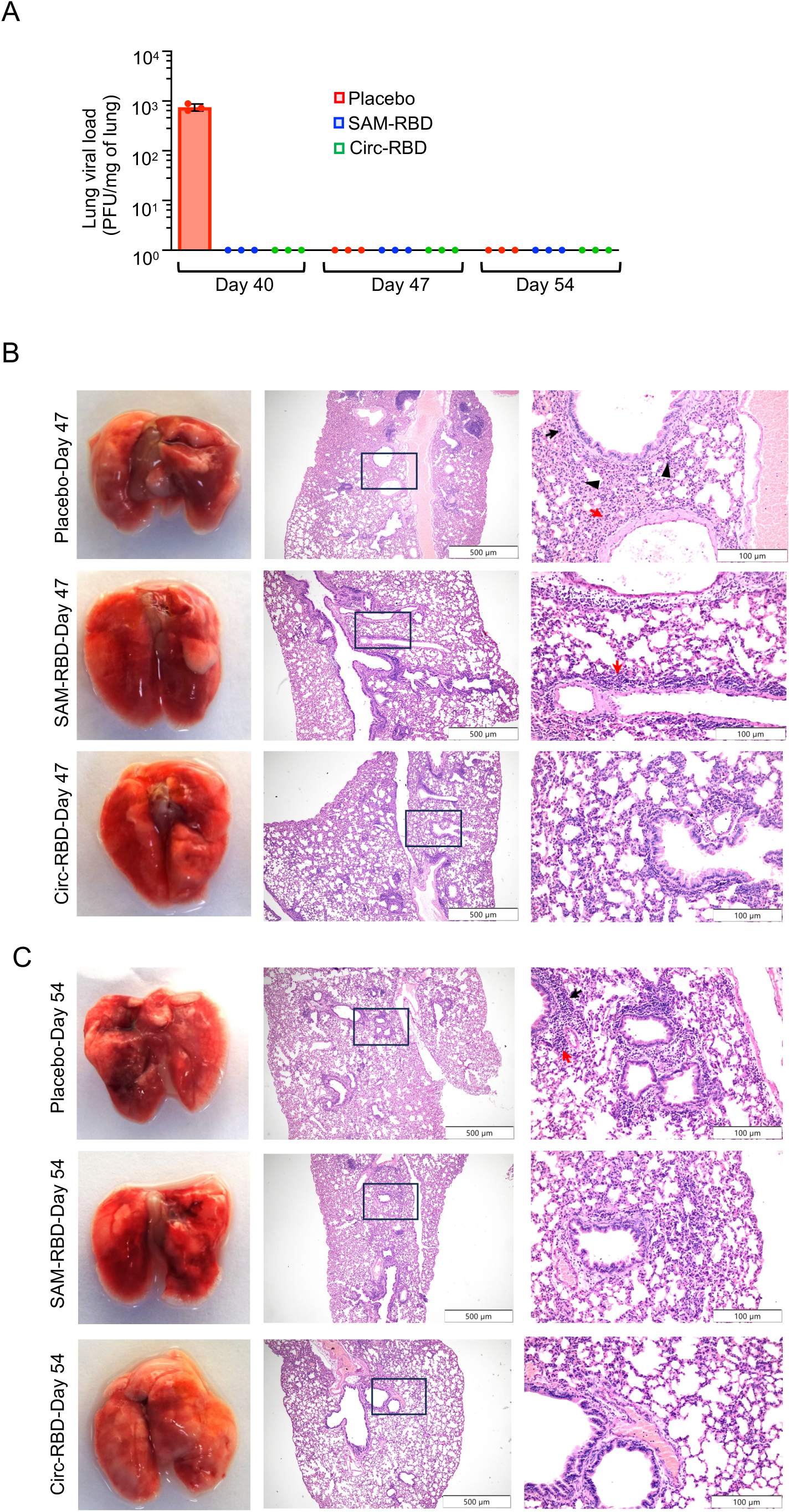
Immunization with the SAM-RBD and Circ-RBD RNA vaccines protect mice from intratracheal SARS-CoV-2 challenge. (A) Measurement of viral load in aliquots of the lung tissue of placebo, SAM-RBD and Circ-RBD RNA vaccine immunized, SARS-CoV-2 (Wuhan) infected mice as indicated in Fig. 3A. Viral load was determined by plaque assay and represented as PFU/mg of lung in samples collected on day 40, 47 and 54. (B) Lung morphology of the placebo, SAM-RBD and Circ-RBD RNA vaccine immunized, SARS-CoV-2 (Wuhan) infected mice, collected on day 47 (left panel). Middle and right panels show representative H&E stained image of lung tissue section from samples shown in left panel. Images were acquired at 40X (middle panel) and 200X magnification (right panel). Black and red arrows indicate peribronchiolar and perialveolar infiltration, respectively, and black arrow-head indicates neutrophils. (C) Lung morphology of the placebo, SAM-RBD and Circ-RBD RNA vaccine immunized, SARS-CoV-2 (Wuhan) infected mice, collected on day 54 (left panel). Middle and right panels show representative H&E stained image of lung tissue section from samples shown in left panel. Images were acquired at 40X (middle panel) and 200X magnification (right panel). Black and red arrows indicate peribronchiolar and perialveolar infiltration, respectively.

